# Multi-Omic Profiling Identifies Conserved Metabolic Pathways Critical for SARS-CoV-2 Variants Infection

**DOI:** 10.1101/2025.06.02.657371

**Authors:** Scotland E. Farley, Jennifer E. Kyle, Helene Jahn, Lisa M. Bramer, Paul D. Piehowski, Athena A. Shepmoes, Brooke LD. Kaiser, Sarai M. Williams, Josie G. Eder, Carsten Schultz, Fikadu G. Tafesse

**Affiliations:** Oregon Health & Science University, Department of Molecular Microbiology and Immunology and; Oregon Health & Science University, Department of Chemical Physiology and Biochemistry, Portland, OR, 97239, USA; Pacific Northwest National Laboratories (PNNL), Biological Sciences Division, Earth and Biological Sciences Directorate, Richland, WA, USA; Pacific Northwest National Laboratories (PNNL), Computational Biology Group, Earth and Biological Sciences Directorate, Richland, WA, USA

## Abstract

The rapid evolution of SARS-CoV-2 has led to the emergence of numerous variants with enhanced transmissibility and immune evasion. Despite widespread vaccination, infections persist, and the mechanisms by which SARS-CoV-2 reprograms host metabolism remain incompletely understood. Here, we investigated whether virus-induced lipid remodeling is conserved across variants and whether changes in lipid abundance correlate with alterations in lipid biosynthetic enzymes. Using global untargeted lipidomics and quantitative proteomics, we analyzed A549-ACE2 cells infected with the Delta (B.1.617.2) or Omicron (B.1.1.529) variants and compared them to cells infected with the ancestral WA1 strain. In parallel, we conducted quantitative proteomics to assess virus-induced changes in the host proteome. Our results reveal that SARS-CoV-2 drives a remarkably consistent pattern of metabolic rewiring at both the lipidomic and proteomic levels across all three variants. We mapped changes in the expression of host metabolic enzymes and compared these to corresponding shifts in lipid abundance. This integrative analysis identified key host proteins involved in virus-mediated lipid remodeling, including fatty acid synthase (FASN), lysosomal acid lipase (LIPA), and ORM1-like protein 2 (ORMDL2). Together, these findings highlight conserved metabolic dependencies of SARS-CoV-2 variants and underscore host lipid metabolism as a potential target for broad-spectrum antiviral strategies.

## Introduction

SARS-CoV-2 has undergone rapid and extensive genetic evolution, giving rise to variants with significant genomic alterations. These emerging variants have caused recurring waves of infection, as each new lineage has differed substantially from both the parent (WA1) strain and earlier variants in their transmissibility and ability to evade adaptive immune responses (1–6). Both the increased transmissibility and immune evasion are driven by mutations in the spike protein, the major antigenic protein localized on the surface of the virion, which mediates attachment and entry into the host cell (7, 8). The last few years have produced a comprehensive body of information about how the spike mutations affect immune responses mounted in response to antigens of previous generations of SARS-CoV-2 virus (1, 2, 4, 5, 9–11). However, how these mutations impact the virus’s ability to manipulate host metabolic pathways remains poorly understood.

While vaccination has significantly reduced COVID-19-related deaths and hospitalizations, a substantial number of individuals continue to suffer from long COVID, an emerging condition with poorly understood pathophysiology (12–14). Growing evidence suggests that dysregulated host metabolism, including lipid metabolism, contributes to long COVID symptoms, and that metabolic modulators may offer therapeutic benefit (15–18). These findings highlight the need to better understand how SARS-CoV-2 alters host lipid metabolism, and whether these effects are conserved across viral variants.

SARS-CoV-2 is an enveloped, positive-sense, single-stranded RNA virus with a large ∼30 kb genome (19). Its genome encodes two large open reading frames (ORFs) that produce 16 non-structural proteins, along with additional ORFs that encode four structural and nine accessory proteins, most of which are essential for viral replication and assembly (20, 21). Paxlovid, which targets the viral protease nsp5 (21), is an example of a successful therapeutic targeting a non-spike viral protein (22). Even more genetically stable than viral proteins are host pathways the virus exploits for replication, providing a strong rationale for host-directed therapies aimed at conserved metabolic dependencies, including lipid pathways (23).

We previously described the rewiring of lipid metabolism in WA1 (original SARS-CoV-2 strain) infection and demonstrated that many aspects of lipid synthesis were essential for viral shedding — the fatty acid synthase inhibitor GSK2194069 had an EC_50_ comparable to the clinical antiviral drug remdesivir(24). Other groups described metabolomic changes in patient samples that mirrored our observations in cell culture (25, 26). However, there have not been updated studies that assess the degree to which this metabolic rewiring is conserved among later SARS-CoV-2 variants, which is critical information that would inform the long-term viability of antiviral strategies that depend on either nonstructural viral proteins or host pathways.

To test the conservation of metabolic rewiring, we compared global nontargeted lipidomics on A549-ACE2 cells infected with the delta variant (B.1.617.2), or the omicron (B.1.1.529) variant to our previous results of global nontargeted lipidomics on A549-ACE2 cells infected with the original WA1 strain, and compared quantitative proteomics of all three virus strains. We mapped changes in expression of lipid biosynthetic enzymes responsible for metabolizing fatty acids, glycerolipids, and sphingolipids, and correlated these findings with changes in the abundance of specific lipid species and classes. Overall, we demonstrate a profound and conserved induction of dihydroceramides and hexosylceramides, and a concurrent decrease in the expression of the sphingolipid regulatory protein ORM1-like protein 2 (ORMDL2), the desaturase that converts dihydroceramides into ceramides. We also observed an increase in *de novo* fatty acid and glycerolipid biosynthesis and a downregulation of enzymes that degrade triacylglycerol (TAG). These results provide a comprehensive map of the remodeling of lipid biosynthetic machinery in SARS-CoV-2-infected cells, suggesting possible candidates for future host-targeted therapeutics that would potentially be less susceptible to the ongoing genetic variability of the spike protein.

## Results

### Variants of concern manipulate the host lipidome in generally conserved patterns

To assess whether virus-induced lipid remodeling is conserved across SARS-CoV-2 variants and whether shifts in lipid abundance correspond to changes in lipid biosynthetic pathways, we conducted parallel global lipidomic and proteomic analyses (Fig 1A). The genomic landscape of SARS-CoV-2 has substantially shifted from the early days of the pandemic (Fig 1B). The first four variants of concern were different enough from the original WA1 strain to drive waves of infection and hospitalization, but the omicron lineage represented an even more dramatic divergence, with an explosion of diversity among spike sequences. To comprehensively compare the changes in lipid composition following infection with variants of SARS-CoV-2, we compared changes in the cellular lipidome and proteome after infection with the original WA1 strain of virus, the delta (B.1.617.2), or the omicron (B.1.1.529/BA1), at an MOI (multiplicity of infection) of 1 for 24 or 48 hours. We selected an MOI that would infect 70-90% of cells without causing cell death (Supplementary Figure 2). While all samples were prepared and analyzed in parallel, the lipidome of WA1-infected A549 cells has been published and thoroughly analyzed previously(24). After infection, we subjected A549-ACE2 cells to a chloroform/methanol extraction, harvesting the chloroform layer to analyze the cellular lipidome and the precipitated protein layer to analyze the cellular proteome. Lipid extracts were analyzed by liquid chromatograph tandem mass spectrometry (LC-MS/MS), using a cocktail of internal lipid standards (Avanti) for quantitative comparison. We identified 443 unique lipid species (Extended Data 1), of which 209 were significantly changed relative to mock in WA1 infection, 279 were significantly changed in delta infection, and 198 were significantly changed in omicron infection (adjusted p < 0.05).

**Figure 1.**
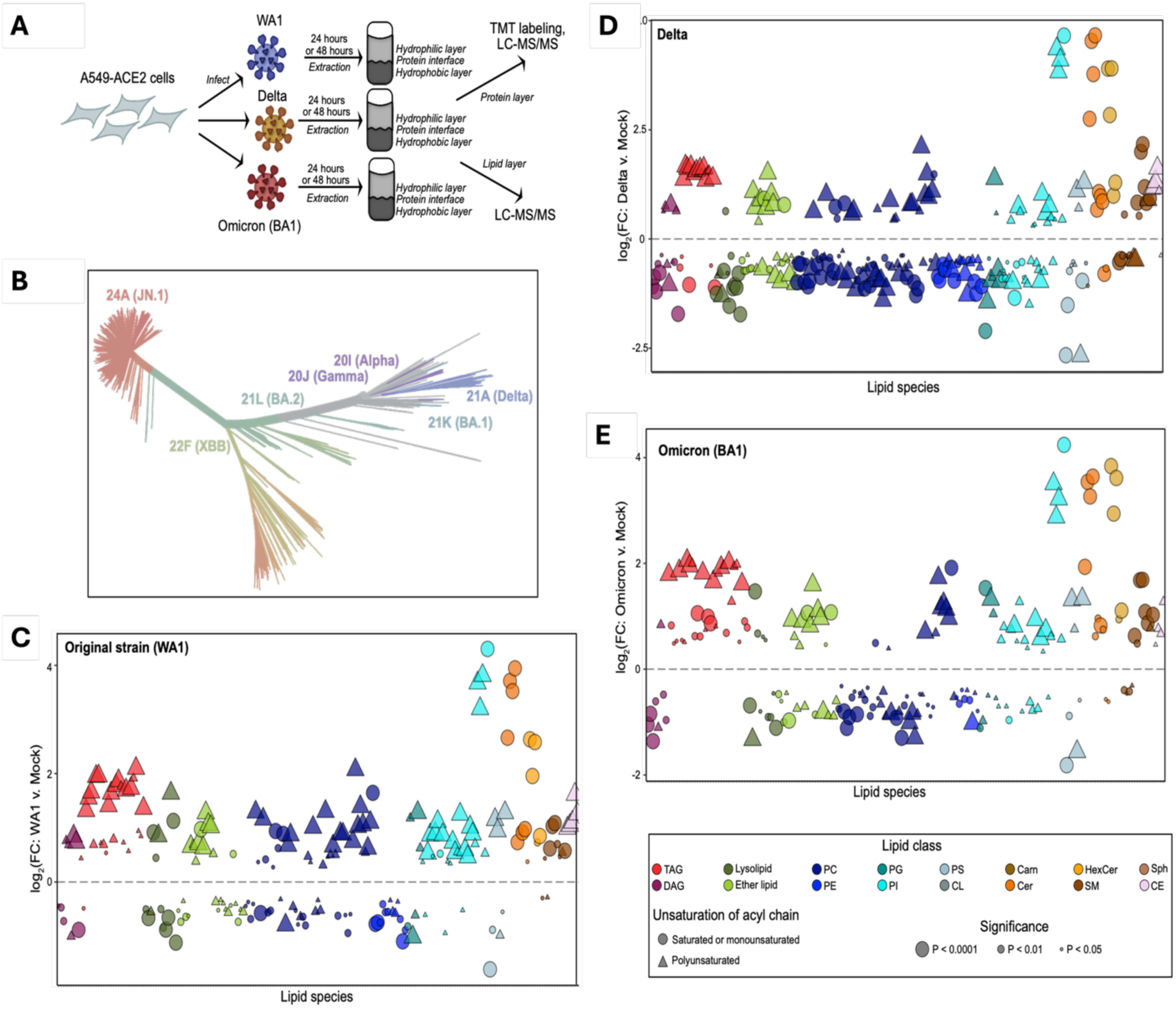
**(A)** Lipidomics study design. (**B**) Phylogenetic map of major variants of SARS-CoV-2 from the NextStrain (27, 28) database, as of June 11, 2024. (**C-E**) Individual lipid species characterized by abundance in SARS-CoV-2 infection (C: original WA1 strain; D: Delta variant; E: omicron BA1 variant) relative to mock in A549-ACE2 cells. Data points are means from 5 biological replicates; each data point represents a lipid species. Lipids were harvested 24 hours after infection. Only significantly changed (p < 0.05, one way ANOVA, with Benjamini-Hochberg adjustment for multiple comparisons) lipids are shown. Log_2_(Fold Change) relative to mock infection is shown on the x-axis. Individual lipid species are colored by the class of lipid that they belong to. DAG = diacylglycerol; TAG = triacylglycerol; PC = phosphatidylcholine; PE = phosphatidylethanolamine; PG = phosphatidylglycerol; PI = phosphatidylinositol; PS = phosphatidylserine; CL = cardiolipin; Cer = ceramide; HexCer = hexosylceramide; SM = sphingomyelin; Carn = acylcarnitine; CE = cholesterol ester.

We observed very similar patterns of lipid abundance changes from mock in all the delta (Fig 1D) and the omicron (Fig 1E) variants, which again closely resemble our previously reported WA1 lipidomics(24). In all three cases, there is an increase in (mostly polyunsaturated) TAG, diverse shifts in a number of phospholipid species, and dramatic increases in ceramides (Cer), especially dihydroceramides (dhCer), and hexosylceramides (HexCer). The same four highly polyunsaturated phosphatidylinositol (PI) species — PI(42:9); PI(20:4/22:6); PI(20:4/20:4); PI(20:3/20:4) — are all dramatically increased in all three viruses, each at least 8-fold enriched over mock infection. This is also true of the top four ceramide species: Cer(d18:0_16:0), Cer(d18:0_22:0), Cer(d18:0_24:0), Cer(d18:0_24:1) and the top three hexosylceramide species: HexCer(d18:0_22:0), HexCer(d18:0_24:0), HexCer(d18:0_24:1).

### Minor differences are observed in lipidomic changes caused by variants of SARS-CoV-2

To look in more detail at the genomic variability of the different viral strains that must underpin the relationship to the host lipidome, we collected the list of mutations from BEI resources (Supplementary Table 1) where each variant diverges from WA1, and counted the number of individual mutations that are present in each viral protein. The dramatic divergence of the omicron lineage is evident — BA.1 has nearly twice the number of spike mutations as any previous variant, and JN.1, the strain of omicron that emerged in early 2024, has twice as many spike mutations as BA.1 (Fig. 2A). Perhaps as striking as the diversity within the spike protein, however, is the relatively modest divergence in the sequences of the nonstructural and accessory proteins; even the JN.1 variant has only a handful of mutations in the non-spike proteins. Thus, we hypothesize that the lineage-driving spike mutations do not play a substantial role in manipulating the intracellular lipid environment.

**Figure 2.**
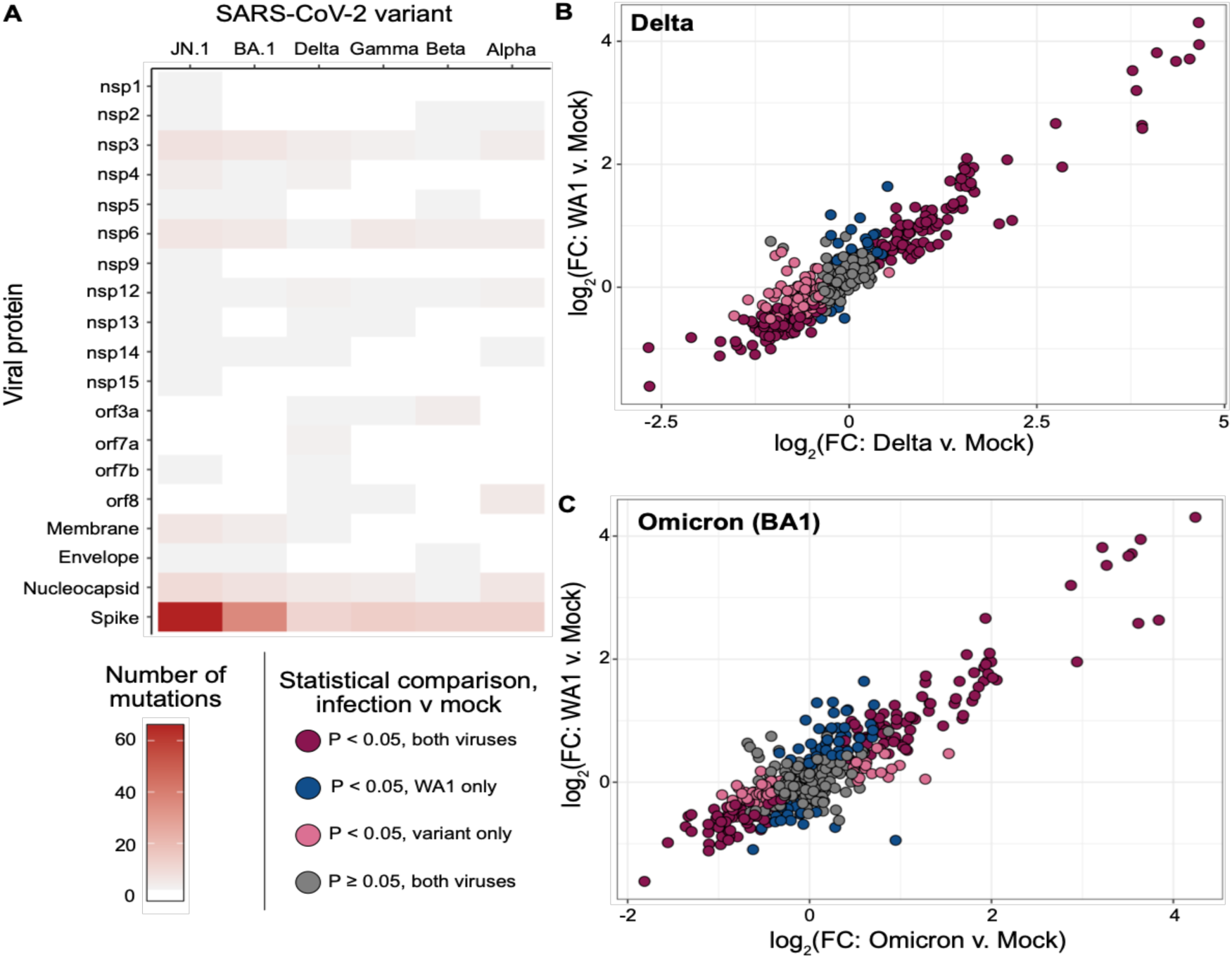
(**A**) Number of mutations in each viral protein, for each indicated variant, relative to the original WA1 strain. Mutation information comes from BEI resources and is detailed in Supplementary Table 1. (**B**) Comparison of the log-2 transformed fold changes from mock of individual lipid species between Delta infection (x axis) or WA1 infection (y axis). All detected lipids are shown and colored by the P-value (one way ANOVA, with Benjamini-Hochberg adjustment for multiple comparisons) for the comparison between that virus and mock infection. (**C**) Comparison of the log-2 transformed fold changes from mock of individual lipid species between Omicron BA1 infection (x axis) or WA1 infection (y axis). All detected lipids are shown and colored by the P-value (one way ANOVA, with Benjamini-Hochberg adjustment for multiple comparisons) for the comparison between that virus and mock infection.

To directly compare the lipidomic changes caused by each variant to the changes observed for the original WA1 strain, the log_2_(FC) of each variant of concern relative to mock was plotted against the log_2_(FC) of WA1 relative to mock (Fig 2B, WA1 versus delta, and Fig 2C, WA1 versus Omicron). Again, we see that all variants induce very similar changes to the host lipidome; in case of differences in lipid abundance, the difference was generally one of magnitude rather than direction, and the differences were rather modest (direct comparisons between variants are shown in Supplementary Figure 3). For example, both Delta and Omicron induced a ∼2-fold increase in three long-chain dhHexCer species, with Delta also elevating two sphingomyelin species to a similar extent. In the neutral lipid family, delta induced a handful of TAG species less strongly than WA1 did, and Omicron induced a handful of TAG species about 1.5-fold more than WA1. The most substantial difference in any lipid species was observed between Omicron and WA1, in the phosphatidylinositol species PI(18:0_20:2), which Omicron induced but WA1 suppressed. Overall, these results indicate a strikingly similar pattern of lipid remodeling among the variants of concern studied here, some subtle differences in the abundance of a subset of lipid species, but with a conservation of the general lipidomic trends.

### Variants of SARS-CoV-2 induce substantial changes in the abundance of lipid-associated proteins

Next, we wanted to assess whether virus-induced changes to the host lipidome could be traced to changes in the expression of host proteins associated with lipid metabolism. To this end, the protein layers of the lipid extractions used for the lipidomics analyses described above were analyzed by LC-MS/MS, using tandem mass tag (TMT) labeling for multiplexing and quantitative comparison.

In total, 8,339 proteins were identified, of which, at 24 hours post-infection, 1,352 were significantly different between WA1 and mock, 1,556 were significantly different between delta and mock, and 949 were significantly different between Omicron and mock. By 48 hours post infection, the proteomic changes after infection were even more pronounced, with 4,423 proteins significantly changed with WA1 infection and 4,235 proteins significantly changed with delta infection (Fig 3A and Extended Data 2). We used principal component analysis to visualize the differences between the different infection groups (Fig 3B) and found that, at 24 hours after infection, each infection condition clustered independently, albeit with some overlap. At 48 hours after infection, interestingly, the distance between the SARS-CoV-2 infected samples and mock samples was more pronounced, while the distance between the different variants (WA1 and delta) was reduced.

**Figure 3.**
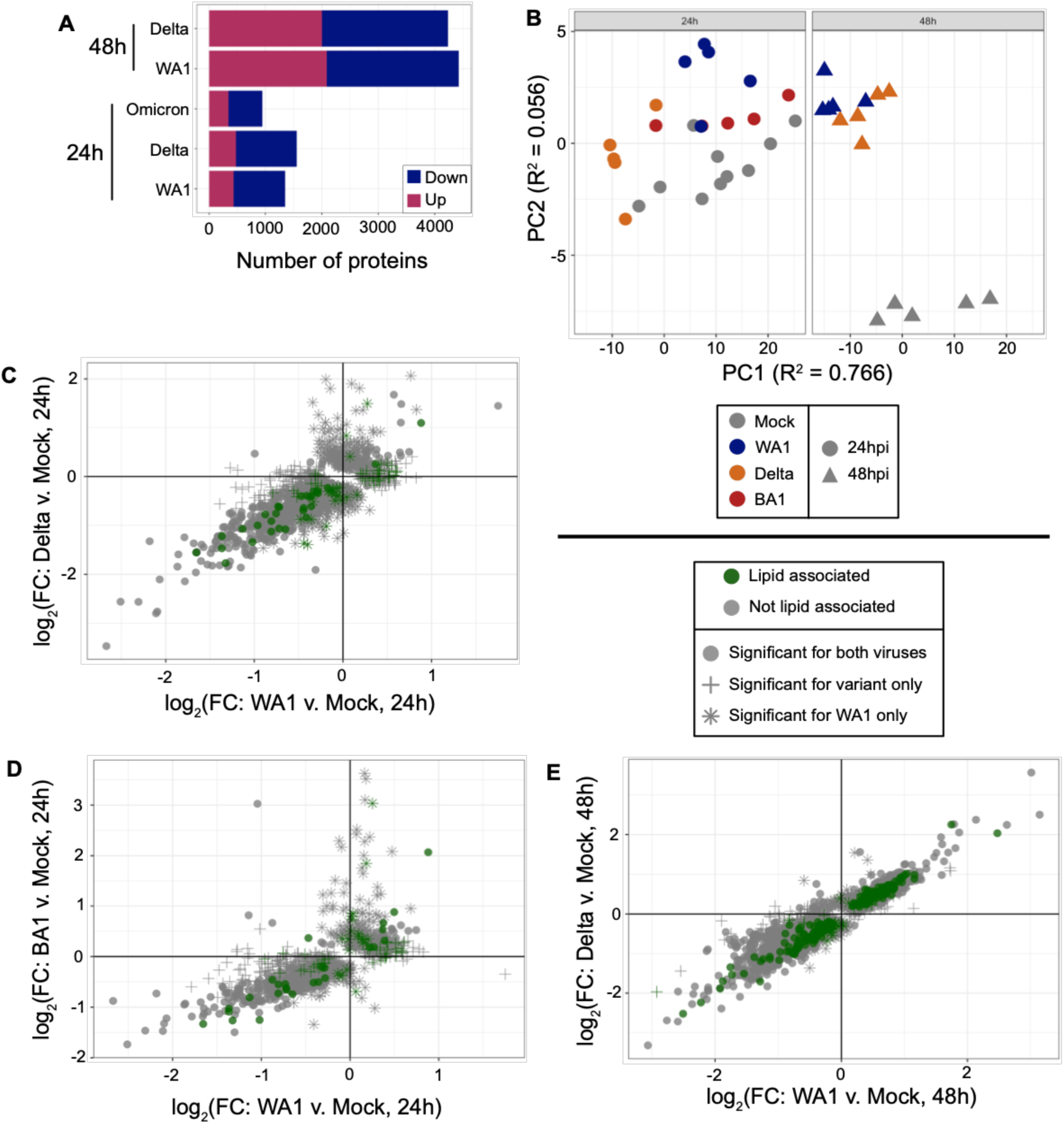
(**A**) Number of proteins whose expression significantly changes relative to mock for each infection and timepoint (one-way ANOVA with Holm adjustment for multiple comparisons). (**B**) Principal Component Analysis (PCA) of global proteomics data. (**C-E**) log_2_FC of WA1 versus mock compared to log_2_FC of a variant of concern versus mock. Proteins that were significantly changed in one or both of the infections are shown.

In order to begin interrogating the relationship between proteomic changes and lipidomic changes, we used the Lipid Maps Protein Database (29), a collection of proteins with annotated association with lipids in public databases (UniProt, EntrezGene, ENZYME, GO, KEGG). These proteins include lipid metabolic enzymes, lipid binding proteins, as well as chaperones, transcription factors, and receptors involved in lipid metabolism and membrane association. We found a substantial minority of lipid-associated proteins altered by WA1 infection: 6.8% of altered proteins at 24 hours and 6.1 % of altered proteins at 48 hours post-infection. Comparing lipid-associated proteins between variants, we find that 38% of the lipid-associated proteins changed in WA1 infection are also changed in delta infection, and 31% of the lipid-associated proteins changed in WA1 infection are also changed in omicron infection. By 48 hours post-infection, the WA1 and delta lipid-associated proteins had further converged, with 88% overlap in their changed lipid-associated proteins. This mirrors an overall convergence in the proteomic changes over time, with about 40% overlap of significantly changed proteins between WA1 and delta at 24 hours post-infection, and 85% overlap at 48 hours post-infection (Fig 3C - 3E).

### Lipid metabolic enzymes across lipid families change in abundance after SARS-CoV-2 infection

Of the lipid-associated proteins that significantly changed in response to WA1 infection, we examined the exact categories of protein more granularly, by manually annotating each protein based on their descriptions in UniProt (Supplementary Table 2). Proteins were categorized as 1) lipid enzymes: enzymes that participate exclusively in lipid biosynthesis; 2) other enzymes: enzymes that are involved in pathways other than lipid biosynthesis; 3) lipid binding: proteins that bind lipids but do not participate in lipid biosynthesis; 4) lipid and other enzymes: enzymes that both participate in lipid biosynthesis and operate on other molecules; 5) protein lipidation: enzymes that post-translationally modify proteins by adding a lipid such as palmitic acid; 6) receptors; 7) transcription factors; 8) chaperones (Fig 4A). The largest category represented were lipid metabolic enzymes, which directly participate in the biosynthesis and turnover of lipids. Because of our interest in the causes of lipidomic shifts following infection, we chose to focus on the lipid metabolic enzymes for a more detailed analysis.

**Figure 4.**
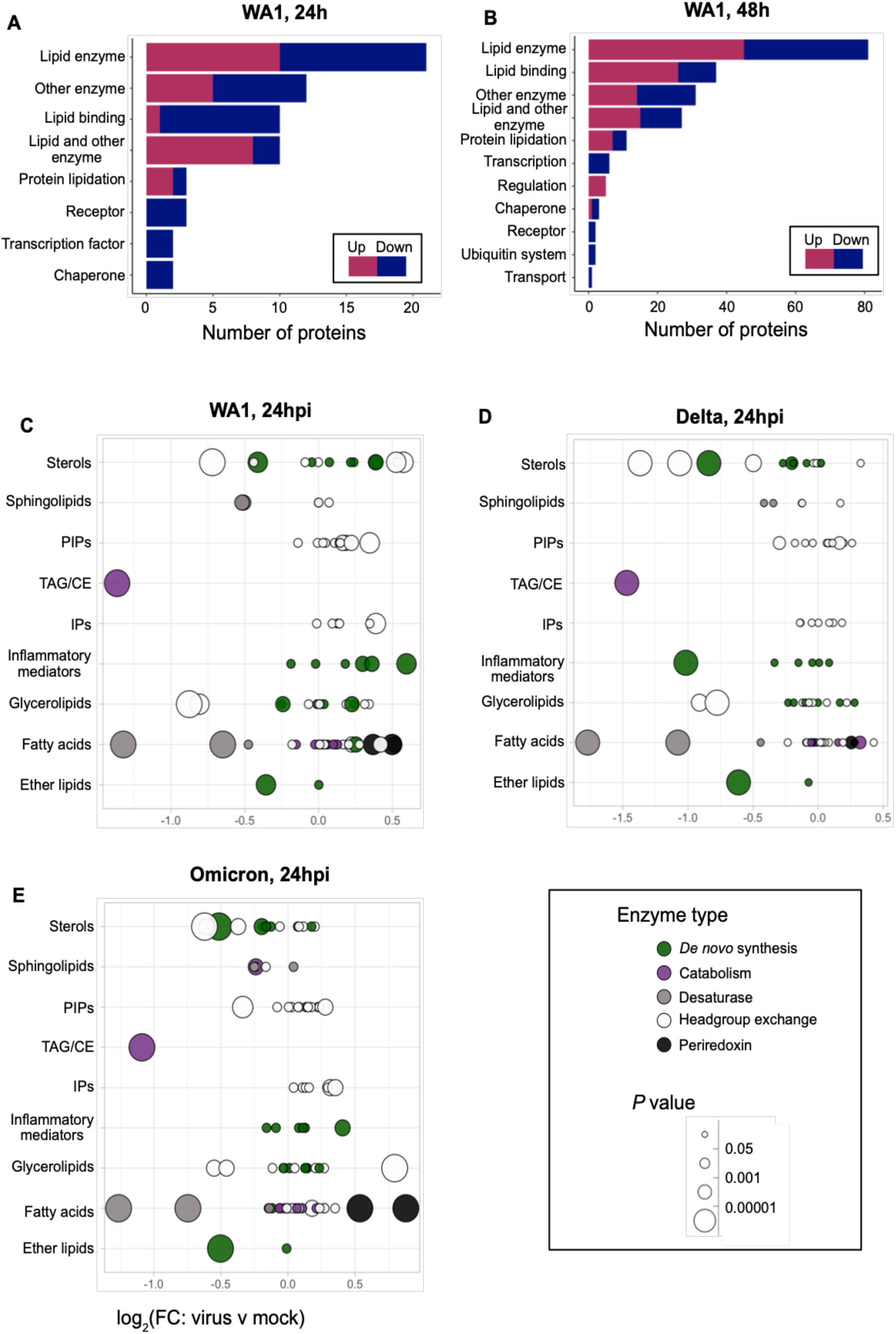
Overview of expression of lipid metabolic enzymes after SARS-CoV-2 infection **(A and B)** Number of lipid associated proteins significantly changed after 24 hours (A) or 48 hours (B) of WA1 infection (P < 0.05, one-way ANOVA with Holm adjustment for multiple comparisons). Pink indicates an upregulated protein; blue indicates a downregulated protein **(C-E)** Types of lipid metabolic enzymes observed, and the classes of lipid they operate on, in WA (C), Delta (D), or Omicron (E) infection. Abbreviations: PIP = phosphatidylinositol phosphate; TAG = triacylglycerol; CE = cholesterol ester; IP = inositol phosphate

In WA1 infection, 21 lipid metabolic enzymes were significantly changed relative to mock at 24 hpi, and 81 were significantly changed at 48 hpi (Fig 4A and 4B). The proteins significantly changed at 48 hpi were for the most part an expansion of those at 24 hpi; only two proteins appeared significant at 24 hpi that did not increase in magnitude at 48 hpi. The enzymes were annotated based on both the type of molecule they operate on (sterols, sphingolipids, phosphatidylinositols, TAG/CE, inflammatory mediators such as prostaglandins and eicosanoids, glycerolipids, fatty acids, and ether lipids) and their function in the biosynthetic pathway (*de novo* synthesis, catabolism, desaturation, headgroup exchange, or peroxide reduction) (Fig 4C-E). While there were some individual differences between variants, the metabolic patterns generally resembled WA1, and the largest and most significant changes (in TAG hydrolysis, fatty acid desaturation and catabolism, prostaglandin and ether lipid synthesis) were widely shared.

### Changes in the expression of key *DE NOVO* biosynthesis enzymes correlate with changes in lipid abundance after infection

Ceramides and TAGs are two of the lipid classes that are changed most substantially after infection. Therefore, we examined in detail whether the expression of enzymes involved in the synthesis and breakdown of ceramides and triacylglycerols were changed after infection. Sphingolipid biosynthesis begins in the endoplasmic reticulum with the condensation of serine and palmitoyl-CoA by the serine palmitoyl transferase (SPT) complex (30).

Ceramide synthases 1-6 (CerS1-6) acylate the sphingoid base sphinganine with a variety of fatty acids, producing dihydroceramides (31), and the canonical 4-*trans* double bond is introduced by the ceramide desaturase DES1(32). A second 14-*cis* double bond can be introduced via fatty acid desaturase 3 (FADS3) (33). Under normal conditions, *de novo* biosynthesis of sphingolipids is kept under tight control, with the ORMDL proteins generally associating with the SPT complex and preventing its activity (34, 35) (Fig. 5A). Sphingolipids — especially ceramides — are among the lipids most strongly induced by all variants of SARS-CoV-2, with some hydroceramide species increased 16-fold over mock infection (Fig. 5B). There is a notable difference between the degree of induction of dihydroceramides (which have a fully saturated sphingoid base backbone) and ceramides (which have a 4-*trans* double bond in the sphingoid base) (Fig. 5B). The top four induced sphingolipids are all dihydroceramides with various acyl chain lengths, between 8- and 16-fold increased in abundance over mock. By comparison, ceramides with the *trans*-double bond are only about doubled over the mock. This pattern, although a bit less dramatic, is repeated in the hexosylceramides.

**Figure 5.**
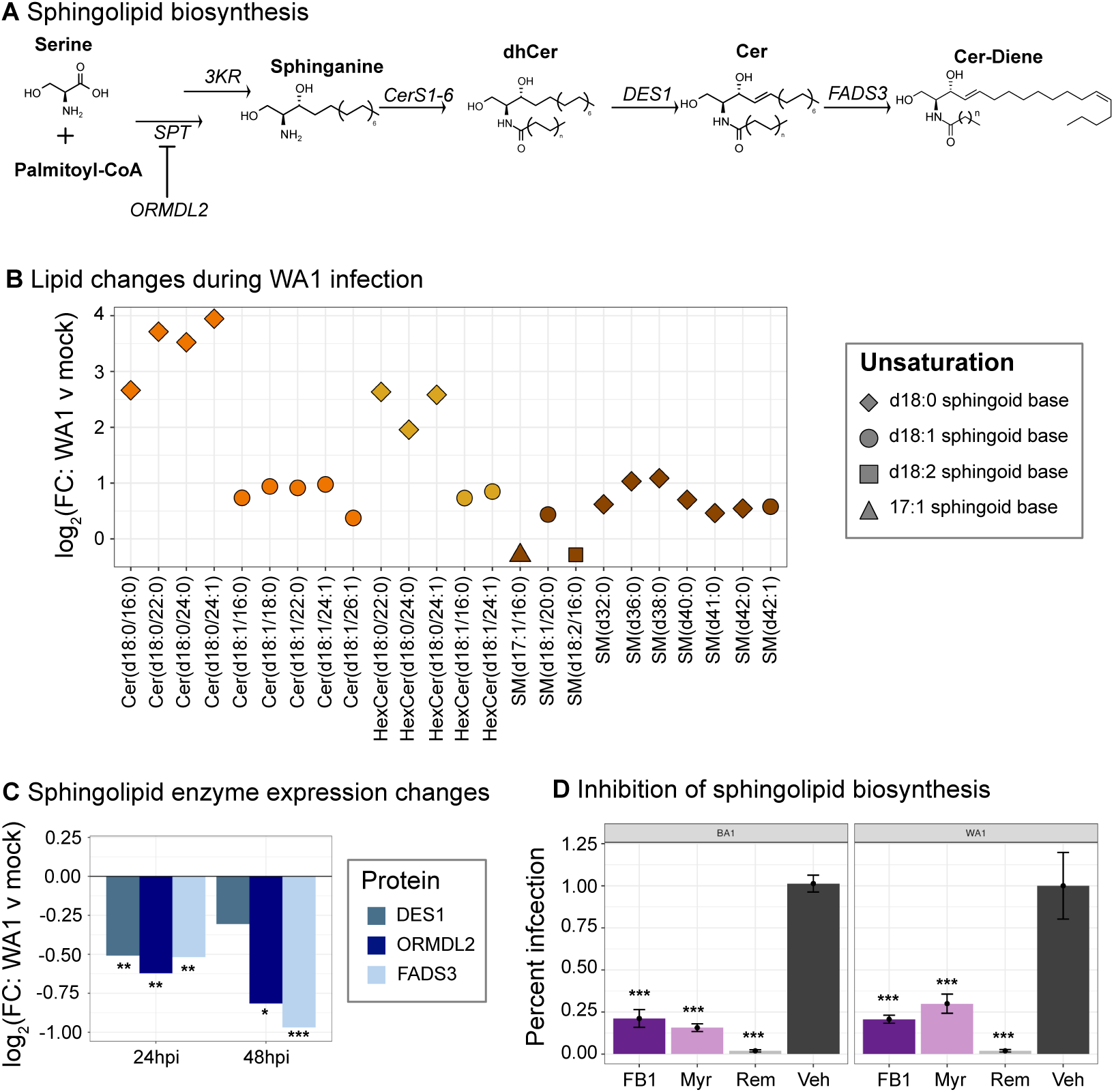
Correlation of lipidomic and proteomic changes in sphingolipid biosynthesis during SARS-CoV-2 infection (**A**) *De novo* sphingolipid biosynthesis. Abbreviations: SPT = serine palmitoyl transferase; ORMDL2 = Orm1-like protein 2; 3KR = 3-keto reductase; CerS = ceramide synthase; dhCer = dihydroceramide; DES1 = ceramide desaturase; Cer = ceramide; FADS3 = fatty acid desaturase 3. (**B**) Sphingolipids that are significantly (P < 0.05, Benjamini-Hochberg adjustment for multiple comparisons) changes during WA1 infection. **(C**) Expression changes for enzymes in the *de novo* sphingolipid biosynthesis pathway (* P < 0.05; ** P < 0.01; *** P < 0.0001; one-way ANOVA with Holm adjustment for multiple comparisons). (**D**) Effect of inhibiting sphingolipid biosynthesis on viral infection, measured by focus forming assay. (* P < 0.05; ** P < 0.01; *** P < 0.0001, one-way ANOVA relative to vehicle control).

Our proteomics data showed significant decreases in the abundance of three enzymes in the *de novo* sphingolipid biosynthesis pathway that are functionally consistent with these lipid changes. The regulatory protein Ormdl2 is significantly reduced, which could allow for higher *de novo* sphingolipid synthesis in SARS-CoV-2 infection. Ceramide desaturase 1 (Des1) is decreased, which could account for the more dramatic increase in dihydroceramides over ceramides. We also observed a dramatic decrease in fatty acid desaturase 3 (FADS3) (Fig 5C). We only observed one 18:2 sphingolipid species (SMd18:2/16:0) in our lipidomics, but it was decreased, consistent with the decrease in abundance of this enzyme.

To assess whether SARS-CoV-2 variants depend on sphingolipid biosynthesis for efficient shedding, we performed a focus forming assay on supernatant from A549-ACE2 cells that had been infected with either WA1 or Omicron, and treated with FB1 (an inhibitor of CerS), myriocin (an inhibitor of SPT), remdesivir (an established inhibitor of coronavirus replication), or DMSO vehicle (Fig 5D). FB1 and myriocin both prevent the production of ceramides and dihydroceramides, and if the presence of these lipid species is required for the virus life cycle, we would expect these inhibitors to prevent viral shedding. We found that both FB1 and myriocin potently reduced shedding at 10 µM, resulting in about 25% infection of the vehicle control. This suggests that the changes in the host lipidome (increase in dihydroceramides) and proteome (decrease in DES1 and ORMDL2) likely represent a necessary manipulation of the host by the virus, rather than being part of the host defense response. The shedding of both the WA1 and Omicron variants was inhibited to similar degrees by inhibitors of sphingolipid biosynthesis, suggesting that dependence on the sphingolipid biosynthesis pathway is conserved from the original WA1 strain to the BA.1 omicron lineage.

The neutral lipids diacylglycerol (DAG) and TAG, as well as cholesterol esters (CE), incorporate fatty acid chains produced by the fatty acid synthase (FASN). TAG is synthesized from DAG by one of two diacylglycerol acetyltransferases (DGAT1/2), and DAG can be produced by the lipolysis of TAG by many lipase enzymes (Fig 6A). In the neutral lipid family, we observed an increase in both TAG and CE species, almost all of which carried at least one polyunsaturated acyl chain (Fig 6B). Our proteomics data showed several proteins in the *de novo* fatty acid pathway whose abundance was significantly changed during SARS-CoV-2 infection, including both AACS (acetoacetyl-CoA synthetase), which acylates coenzyme A in order to initiate fatty acid synthesis(^36^), and FASN itself. There was also a decrease in LICH (lysosomal acid lipase)(^37^), a lysosomal hydrolase that degrades both TAG and CE (Fig 6C). Taken together, these proteomic changes support a lipid metabolic environment with both increased synthesis of fatty acids and reduced lysosomal degradation of storage-neutral lipids.

**Figure 6.**
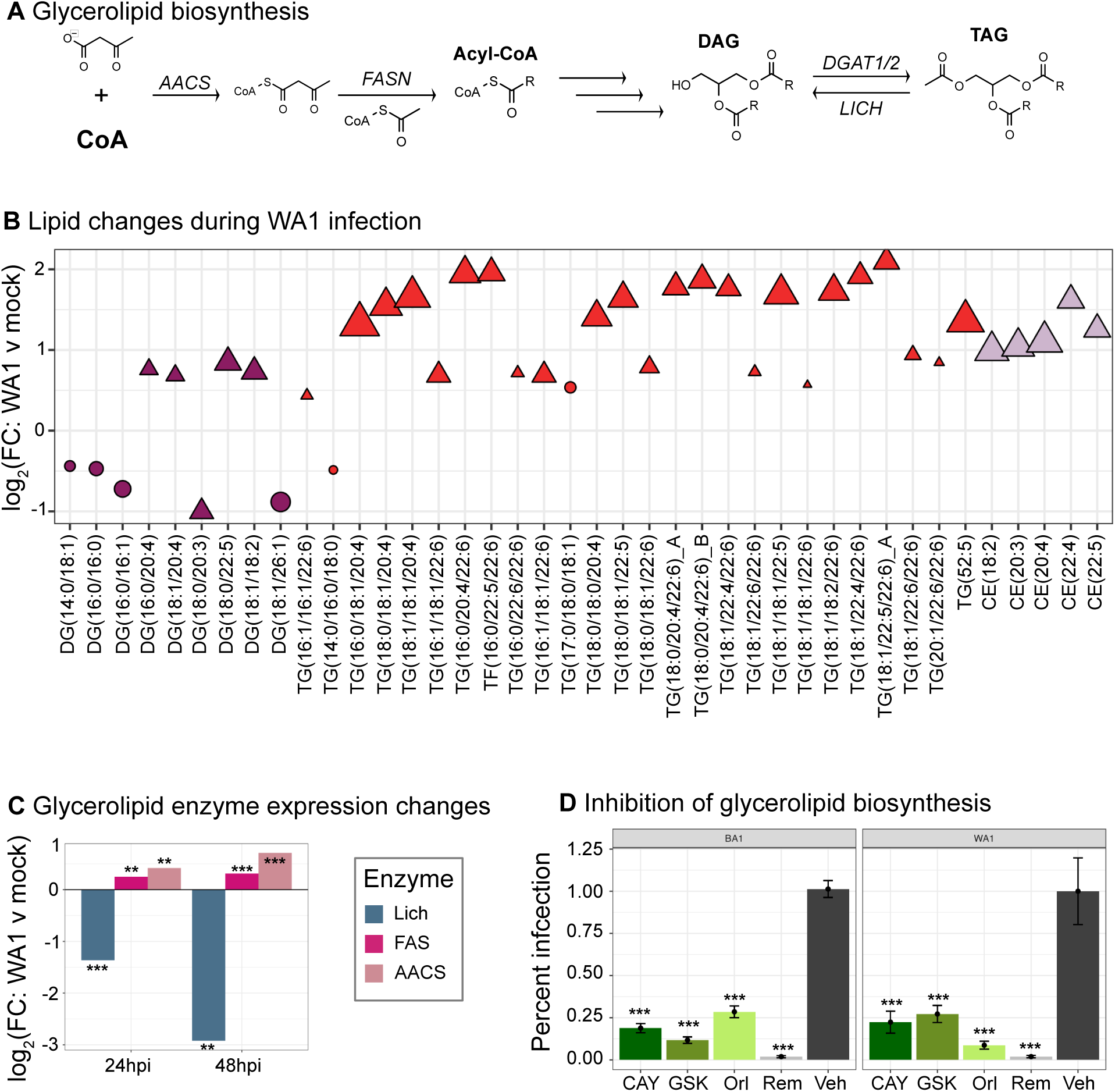
Correlation of lipidomic and proteomic changes in neutral lipid biosynthesis during SARS-CoV-2 infection (**A**) Neutral lipid biosynthesis. Abbreviations: CoA = coenzyme A; AACS = acetyl-CoA synthetase; FASN = fatty acid synthase; DAG = diacylglycerol; DGAT = diacylglycerol acetyltransferase; LICH = lipase A, lysosomal acid type; TAG = triacylglycerol (**B**) Neutral that are significantly (P < 0.05, Benjamini-Hochberg adjustment for multiple comparisons) changes during WA1 infection. **(C**) Expression changes for enzymes in the *de novo* neutral lipid biosynthesis pathway (* P < 0.05; ** P < 0.01; *** P < 0.0001; one-way ANOVA with Holm adjustment for multiple comparisons). (**D**) Effect of inhibiting glycerolipid biosynthesis on viral infection, measured by focus forming assay. (* P < 0.05; ** P < 0.01; *** P < 0.0001, one-way ANOVA relative to vehicle control for each virus).

To further validate our lipidomic results, we inhibited the biosynthesis of key lipid-modifying enzymes using small-molecule inhibitors targeting glycerolipid metabolism during WA1 and Omicron infections (Fig 6D). Viral shedding was quantified by focus forming assay in inhibitor-treated cells and compared to vehicle controls. We used three distinct inhibitors: CAY10499, a broad lipase inhibitor that prevents TAG breakdown; GSK2194069, a selective inhibitor of fatty acid synthase (FASN); and orlistat, which inhibits both TAG lipases and FASN. Blocking these lipid metabolic pathways significantly reduced viral shedding in both WA1 and Omicron infections (Fig 6D). This once again suggests that the changes in protein expression that we observe here — an increase in *de novo* fatty acid synthesis enzymes AACS and FASN, and an increase in the lipolytic enzyme LICH — are part of essential viral reprogramming of host lipid metabolism, rather than being part of the host defense response. These results further underscore the fact that variants of SARS-CoV-2 appear to require enhanced flux through the neutral lipid pathway and increased *de novo* fatty acid synthesis, consistent with our previous findings for WA1 infection(24).

## Discussion

In this study, we integrate global lipidomics and global proteomics of three strains of SARS-CoV-2 to interrogate the ways variants of this virus manipulate host lipid biosynthesis. We describe a generally conserved alteration of the host lipidome by the original WA1 strain, the delta (B. 1.617.2) and omicron (BA.1) strains. These viruses generally suppress phosphatidylethanolamines, DAG, and lysolipids, while inducing phosphatidylinositols, dihydroceramides, hexosylceramides, and TAG. While there are some statistically significant differences between WA1 and either delta or Omicron, these are largely differences in the degree to which the virus induces or suppresses a specific lipid species, rather than a difference in the direction that the lipid is altered. Changes in the abundance of lipid-associated proteins were less conserved than changes in the abundance of lipids themselves, with only about 40% overlap between WA1 and delta 24 hours after infection. However, changes in many of the lipid metabolic enzymes altered by infection were shared among the variants, especially the most dramatically altered enzymes. We describe a profound and conserved induction of dihydroceramides and hexosylceramides, and a concurrent decrease in the expression of the sphingolipid regulatory protein ORMDL2, and DES1, the desaturase that converts dihydroceramides into ceramides. We also observed an increase in *de novo* fatty acid and glycerolipid biosynthesis via AACS and FASN, and a downregulation of the TAG and CE lipase LICH.

The mechanisms by which lipid levels are dynamically manipulated are complex, both in normal eukaryotic biology, and during viral infection. They are not under direct genetic control — while the expression and abundance of lipid biosynthetic enzymes is certainly a prerequisite for producing an individual lipid, other factors are at play as well. Relocalizing lipid enzymes to different subcellular compartments, to give them more (or less) access to their substrates, is a strategy by which a high concentration of a given lipid can be transiently created in a specific location, without changing the abundance of that enzyme, as in the case of membrane recruitment of phosphatidylinositol-4 kinase B (PI4KB) by ACBD3 to produce PI4P (35). Repressing or increasing the activity of an enzyme, again without changing its expression, is another very common strategy; this is the principle by which ORMDL inhibits *de novo* sphingolipid biosynthesis^24^. In the context of viral infection, for example, dengue virus recruits host fatty acid synthase to the sites of virus replication and directly increases its activity through interaction with the viral NS3 protein (36). The changes in abundance of various lipid biosynthetic enzymes that we describe here is certainly only part of the picture of how SARS-CoV-2 manipulates the host lipidome, but it is a critical layer of information upon which to build our understanding of other mechanisms that may be unrelated to control of enzyme expression.

Understanding and managing long COVID remains a major gap in our treatment of this virus. Our study reinforces the idea, already proposed by others, that altered mitochondrial dynamics and metabolism might be at the heart of long-term metabolic damage that seems to mediate at least some of the symptoms involved in long COVID. Many of the pathways we describe herein, from fatty acid synthesis and beta oxidation, to lipolysis of TAG esters to liberate free fatty acids, to sphingolipid metabolism, are either directly located in the mitochondria or influence mitochondrial function. The alteration of many of these lipid species and the enzymes in their metabolic pathways are strikingly conserved among variants, suggesting that they may be part of a fundamental aspect of the SARS-CoV-2 life cycle. Although these are interesting observations in isolation, the relationship between altered lipid metabolism and mitochondrial function in SARS-CoV-2 infection requires more study to determine what a causal relationship might be, and how it is required for other aspects of the viral life cycle. This information will provide both valuable insight into the viral pathogenesis, and potentially very valuable therapeutic avenues.

Here, we demonstrate that, despite their genetic diversity, the WA1, delta, and BA1 variants of SARS-CoV-2 have similar overall effects on the abundance of host lipids, and that while these viruses induce diverse responses in the expression of host proteins, they induce and repress a similar subset of lipid biosynthetic enzymes. This suggests that therapies developed against those non-spike proteins of the virus that mediate replication and manipulate the subcellular environment could have more long-term efficacy across more diverse SARS-CoV-2 variants. Furthermore, this suggests that targeted therapies directed against the host’s metabolic machinery could be a robust antiviral strategy that has a high likelihood of remaining effective against future variants. Many of the pathways described here do not have specific inhibitors against the enzymes involved, and where inhibitors exist, they are far from being optimal clinical candidates. This underscores the need for further development of tools to study lipid turnover, both to inform our understanding of basic lipid biosynthesis and to advance our ability to combat new and existing viruses.

## Acknowledgments

This work is supported by NIH NIAID grant R01AI141549 (F. G. T) and NIH NIGMS RO01 R01 GM127631 (C. S.). Lipidomics and proteomics analyses were supported by Laboratory Directed Research and Development Program (J. E. K., B.LD.K) at Pacific Northwest National Laboratory. The mass spectrometry work was performed at the Environmental Molecular Sciences Laboratory, a U.S. Department of Energy National Scientific User Facility located at the Pacific Northwest National Laboratory operated under contract DE-AC05-76RL01830.

## Materials and Methods

### Cell lines

A549-ACE2 cells were obtained from BEI resources (identifier # NR-53821); Vero E6 cells were obtained from ATCC (identifier # CRL-1586).

### Viral strains

SARS-CoV-2 viral strains were obtained from BEI resources and propagated in Vero E6 cells. Strains:

- “WA1”: isolate USA-WA1/2020: Identifier #NR-5228
- “Delta”: isolate hCoV-19/USA/PHC658/2021: Identifier # NR-55611
- “Omicron”: isolate hCoV-19/USA/MD-HP20874/2021: Identifier # NR-56461

### Reagents

EquiSPLASH lipidomics internal standard was obtained from Avanti Polar Lipids (Identifier # 330731). HPLC-grade solvents were used for lipid extractions, obtained from Fisher Chemical (Chloroform, identifier # C607-4; water, identifier # W5-4; methanol, identifier # A454-1). Anti-N antibody was obtained from Sino Biological (Catalog MA14AP1502).

### Cell culture

Unless otherwise stated, cells were maintained at all times in standard tissue culture-treated vessels in media supplemented with 10% FBS, 1% nonessential amino acids, and 1% penicillin-streptomycin at 37 °C and 5% CO_2_. Vero-E6 cells were cultured in DMEM media, while A549-ACE2 cells were cultured in F12-K media.

### SARS-CoV-2 growth and titration

All SARS-CoV-2 isolates were obtained from BEI resources, as described above. To propagate each virus strain, sub-confluent monolayers of Vero E6 cells were inoculated with the clinical isolates (MOI < 0.01) and grown for 72 hours, at which time significant cytopathic effect was observed for all strains. Culture supernatants were removed, centrifuged 10 min at 1,000 x g, and stored in aliquots at −80°C. To determine titer, focus forming assays were performed on the culture supernatant (assay described in detail below).

### Analysis of variant infection efficiency

A549-ACE2 cells were seeded in glass-bottomed 24-well plates and grown to 75% confluency. Cells were inoculated with each virus (WA1, delta, or BA1), either using pure virus (MOI ∼ 2.5) or an MOI of 1, 0.2, or 0.04, for 1 hour at 37 °C in 2% FBS Opti-MEM, rocking gently every 15 minutes. After 1 hour, infection media was removed and replaced with normal 10% DMEM. Plates were fixed with 4% PFA, blocked and permeabilized using 2% FBS and 0.1% Triton-X-100 in PBS. Infection was visualized by immunostaining with an anti-N primary antibody (1:500 in blocking buffer, overnight at 4 °C), and an Alexa 555 anti-rabbit secondary antibody. DAPI was used to visualize nuclei. An MOI of 1 was chosen for subsequent lipidomic and proteomic experiments.

### Lipidomics and proteomics — Infection

A549-ACE2 cells were seeded to 70% cell density (about 1.5 x 10^6 cells per 10cm dish). Cells were then inoculated with each virus strain (MOI = 1) for 1 hour at 37°C in 2% FBS Opti-MEM, rocking gently every 15 minutes. Mock infections were performed using 2% FBS Opti-MEM for one hour. After 1 hour, infection media was removed and replaced with normal 10% DMEM. Cellular lipids were extracted 24 hour after infection for lipidomics, and 24 or 48 hours after infection for proteomics. Five biological replicates were infected for each condition.

### Lipidomics and Proteomics—sample preparation

Total lipid extraction was performed using a modified Bligh and Dyer method(38), which also enabled simultaneous extraction of total protein from the same samples. Specifically, we adapted the metabolite, protein, and lipid extraction (MPLEx) protocol, a solvent-based method widely used for mass spectrometry-based multi-omics analysis (39). In this approach, the organic phase was collected for lipid analysis, while the aqueous phase was retained for protein extraction.

Briefly, infected and uninfected control cells were harvested from tissue culture dishes by scraping and washed three times with cold PBS. Cell pellets were resuspended in a 2:1:0.75 mixture of chloroform:methanol:water, and 10 µL of an internal standard cocktail (Avanti EquiSPLASH) was added. Samples were incubated at 4 °C for 1 hour, followed by centrifugation at 3,000 × g for 10 minutes to separate the phases. The chloroform layer was transferred to a fresh tube. An additional 2 mL of chloroform was added to the aqueous phase, mixed, incubated again at 4 °C for 1 hour, and centrifuged to recover the remaining lipids. The resulting organic phases were combined and dried under a gentle stream of nitrogen. The aqueous phase was centrifuged at 16,000 × g for 10 minutes to pellet precipitated proteins. The supernatant was discarded, and the protein pellets were retained. Both dried lipid extracts and protein pellets were stored at −80 °C and shipped on dry ice to the Pacific Northwest National Laboratory for LC-MS/MS analysis.

### Lipidomics — LC-MS/MS analysis and lipid identification

LC-MS/MS parameters and lipid identifications were performed as previously described(24, 40). Lipid extracts were analyzed using a Waters Acquity UPLC H-Class system coupled to a Velos-ETD trap mass spectrometer. Dried lipid extracts were reconstituted in 10 µL of chloroform and 540 µL of methanol. A 10 µL aliquot was injected onto a Waters CSH reverse-phase column (3.0 mm × 150 mm, 1.7 µm particle size), and lipids were separated using a 34-minute gradient. The mobile phases consisted of: (A) acetonitrile/water (40:60) with 10 mM ammonium acetate, and (B) acetonitrile/isopropanol (10:90) with 10 mM ammonium acetate. The flow rate was maintained at 250 µL/min. Data were acquired in both positive and negative ionization modes. Fragmentation was achieved using both higher-energy collision dissociation (HCD) and collision-induced dissociation (CID) to maximize lipid identification.

For quality control, MS calibrations were performed prior to all analyses, per ionization mode, and at least once a week, or if mass accuracy > 5 ppm. Solvent blanks of 90:10 ChCl_3_:MeOH were injected prior to and throughout the analysis to monitor for potential contamination. A QC sample, Brain Total Lipid Extract (Avanti Polar Lipids, Inc) was injected every 14 samples to monitor instrument performance including sensitivity, mass accuracy, and retention time drift.

Lipid species were identified using established fragment ion signatures. Raw LC-MS/MS data files were processed with LIQUID (40), followed by manual validation of each identification. Fragmentation spectra were examined for diagnostic ions and acyl chain-specific fragment ions. Additional validation criteria included the precursor ion isotopic profile, mass measurement error, extracted ion chromatogram, and retention time for each lipid species. To enable accurate quantification, a reference lipid database was constructed from the MS/MS-identified lipids. Features from each sample were aligned to this reference based on mass-to-charge ratio (m/z) and retention time using MZmine (40). All aligned features were manually verified, and peak apex intensity values were extracted for statistical analysis.

### Lipidomics — QC, normalization, and statistical comparison methods

Lipidomics data were collected in positive and negative ionization modes and analyzed using R v 4.4.3 and the *pmartR* package (41). Data were log2 transformed. A robust Mahalanobis distance based on lipid abundance vectors (rMd-PAV) was calculated to identify potential sample outliers in the data (42). Two samples were identified as potential outliers (p-value < 0.0001), were verified as outliers based on principal component analysis (PCA), and were removed from the dataset. Deuterated lipids representing various lipid classes (EquiSPLASH, Avanti) were spiked into the samples and used as internal standards (ISs). The pooled coefficient of variation (CV) for each IS was calculated, taking into account strain and sample batch, to assess the consistency of ISs. All ISs were found to have a CV value less than 30%, except 18:1(d7) Chol Ester from the positive mode which had a CV greater than 80%. The median value of ISs for each sample and ionization mode was calculated and used to normalize the data, after excluding 18:1(d7) Chol Ester, via (log2(abundance/median IS abundance)). A one-way analysis of variance (ANOVA) was run on each lipid, and hypothesis tests for all post-hoc pairwise comparisons between virus strains were conducted. A Tukey multiple test correction (43) was used to adjusted for multiple comparisons within each lipid, and a false discovery rate adjustment was made using BH adjustment(44).

### Proteomics — LC-MS/MS analysis

Isolates were reduced with 10 mM dithiothreitol (DTT) for 30 minutes at 37 °C, then alkylated with 40 mM iodoacetamide for 45 minutes at room temperature in the dark. Proteins were digested for 2 hours at 25 °C using sequencing-grade trypsin (1:20 enzyme : protein ratio). Peptides were desalted on a C18 trap column. Desalted peptides were labeled with a 16-plex TMT reagent (15 samples per TMT-plex, with the 16th channel used for a pooled reference sample to compare between plexes), according to the manufacturer’s instructions (ThermoFisher Scientific). Peptides labeled with different TMT reagents were then mixed, dried using a Speed-Vac, reconstituted in 3% acetonitrile, 0.1% formic acid and desalted on tC18 SPE columns. Peptides were separated on a Waters NanoAquity UPLC system with a custom packed C18 column (70cm x 75 µm i.d., Phenomenex Jupiter, 3 µm particle size, 300 Å pore size) coupled to an Q-Exactive HF-X mass spetrometer (Thermo Fisher Scientific), using a water (solvent A) and acetonitrile (solvent B) gradient, both containing 0.1% formic acid: 1–8% B in 2 min, 8–12% B in 18 min, 12–30% B in 55 min, 30–45% B in 22 min, 45–95% B in 3 min, hold for 5 min in 95% B and 99–1% B in 10 min. Peptides were analyzed by nanoelectrospray ionization. Full mass scans were collected from 300 to 1800 m/z at a resolution of 60,000 at 400 m/z. Tandem mass spectra were collected in data-dependent acquisition of the top 12 most intense parent ions using high-energy collision induced dissociation (HCD) fragmentation (0.7 m/z isolation width, 30% normalized collision energy; 30,000 resolution at 400 m/z), before being dynamically excluded for 45 s. Peptides were identified with MaxQuant software (v.1.6.5.0). Resultant Thermo RAW files were processed using *mzRefinery* to correct for mass calibration errors, and then spectra were searched with MS-GF+ v9881(45–47) to match against the UniProt reference protein sequence database from April 2020 (20,352 entries), combined with common contaminants (e.g., trypsin, keratin). A partially tryptic search was used with a ± 20 ppm parent ion mass tolerance. A reversed sequence decoy database approach was used for false discovery rate calculation. MS-GF+ considered static carbamidomethylation (+57.0215 Da) on Cys residues and TMT modification (+229.1629 Da) on the peptide N terminus and Lys residues, and dynamic oxidation (+15.9949 Da) on Met residues. The resulting peptide identifications were filtered to a 1% false discovery rate at the unique peptide level.

### Proteomics — QC, normalization and statistical comparison methods

All analyses were conducted analyzed using R v 4.4.3 and the *pmartR* package (41). Internal reference scaling (IRS) was done to normalize each sample’s peptide abundance profile to the respective TMT plex’s reference pool sample (48). Peptides without enough data to conduct a statistical test were filtered from the data (49). A robust Mahalanobis distance based on lipid abundance vectors (rMd-PAV) was calculated to identify potential sample outliers in the data (42) no sample outliers were identified in the data. Data were normalized via global median centering, and proteins were quantified using a reference-based rollup, rollup (50). For each protein, a oneway analysis of variance (ANOVA) was run on each lipid, and hypothesis tests for all post-hoc pairwise comparisons between virus strains were conducted. A Tukey multiple test correction (43) was used to adjusted for multiple comparisons within each lipid, and a false discovery rate adjustment was made using BH adjustment (44).

### Proteomics — identification of lipid-associated proteins

The Lipid Maps Protein Database (LMPD) (29) was downloaded 11/16/22, and proteins in the infection dataset that were found in the LMPD were designated “lipid associated”. These lipid-associated proteins were further annotated as “lipid enzyme” “lipid binding” “other enzyme” “lipid and other enzyme” “protein lipidation” “receptor” “transcription factor” “chaperone” “ubiquitin system” or “transport” based on their descriptions in UniProt (see Supplementary Table 2).

### Focus forming assay with inhibitors of lipid biosynthesis

A549-ACE2 cells were seeded in 96-well plates at a density of 4,000 cells per well and treated overnight with each inhibitor at a concentration of 10 µM, prior to infection with either WA1 or omicron (see strain information above) at an MOI of 0.1. The infection was continued for 48 h. At 48 hours, supernatant was used to infect a focus forming assay with Vero E6 cells, as described previously(24). Briefly, a confluent monolayer of Vero E6 cells was infected with supernatant from infected A549-ACE2 cells, at several 10-fold dilutions, for 1 hour at 37 °C. After 1 hour, overlay media was added and plates were incubated at 37 °C for 24 hours. Overlay media was then removed, plates were fixed with 4% for at least one hour. Plates were stained with a primary antibody (alpaca anti-SARS-CoV-2 serum; 1:5,000 for WA1 infection, 1:1,000 for omicron infection), then a secondary antibody (anti-llama HRP, goat IgG), and visualized using TrueBlue peroxidase substrate.

## Supporting information

**Supporting information: summary**

Supplementary Figure 1: Pre-processing of lipidomics dataset (related to Figures 1 and 2)

Supplementary Figure 2: infection efficiency of SARS-CoV-2 variants in A549-ACE2 cells (related to all Figures)

Supplementary Figure 3: Statistical comparison of lipidomics results between variants (related to Figures 1 and 2)

Supplementary Table 1: Point mutations in the genome of each SARS-CoV-2 variant (related to Figure 2)

Supplementary Table 2: lipid-associated proteins that are statistically changed by WA1 infection (24 hours after infection), with annotated functions (related to Figures 3 and 4)

Extended Data 1: fold changes and p-values for all lipids that are significantly changed by the infection of any variant

Extended Data 2: fold changes and p-values for all proteins that are significantly changed by the infection of any variant

**Supplementary Figure 1.**
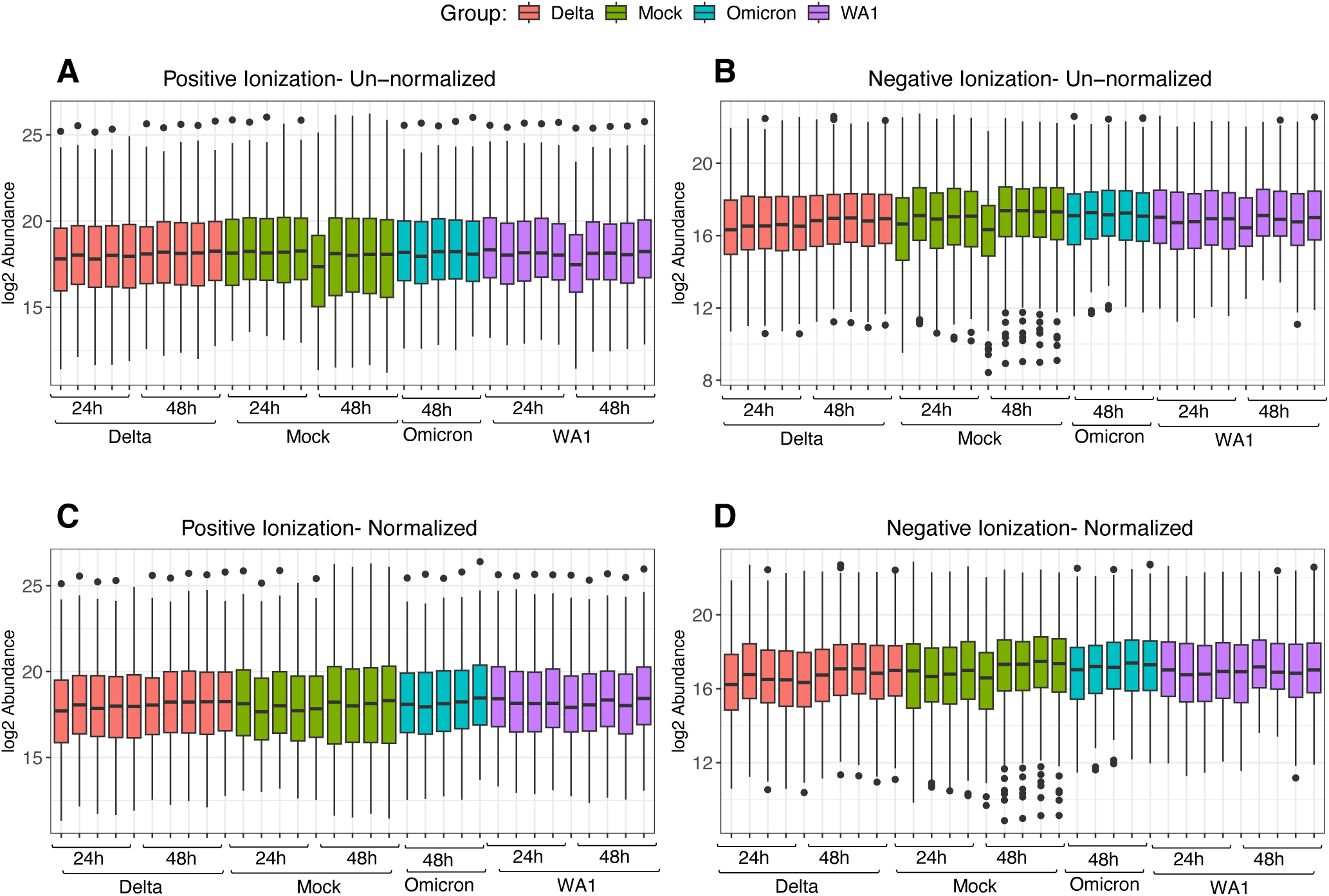
Pre-processing of lipidomic data sets from mock and SARS-CoV-2 variant infected samples. Log_2_ abundance of lipids across full mass spectrometry before (**A, B**) and after (**C, D**) normalization. Boxplots represent median (black line) and 25^th^ to 75^th^ percentile of data (lower and upper bounds of boxes, respectively); whiskers extend to maximum and minimum samples within 1.5 times the interquartile range.

**Supplementary Figure 2.**
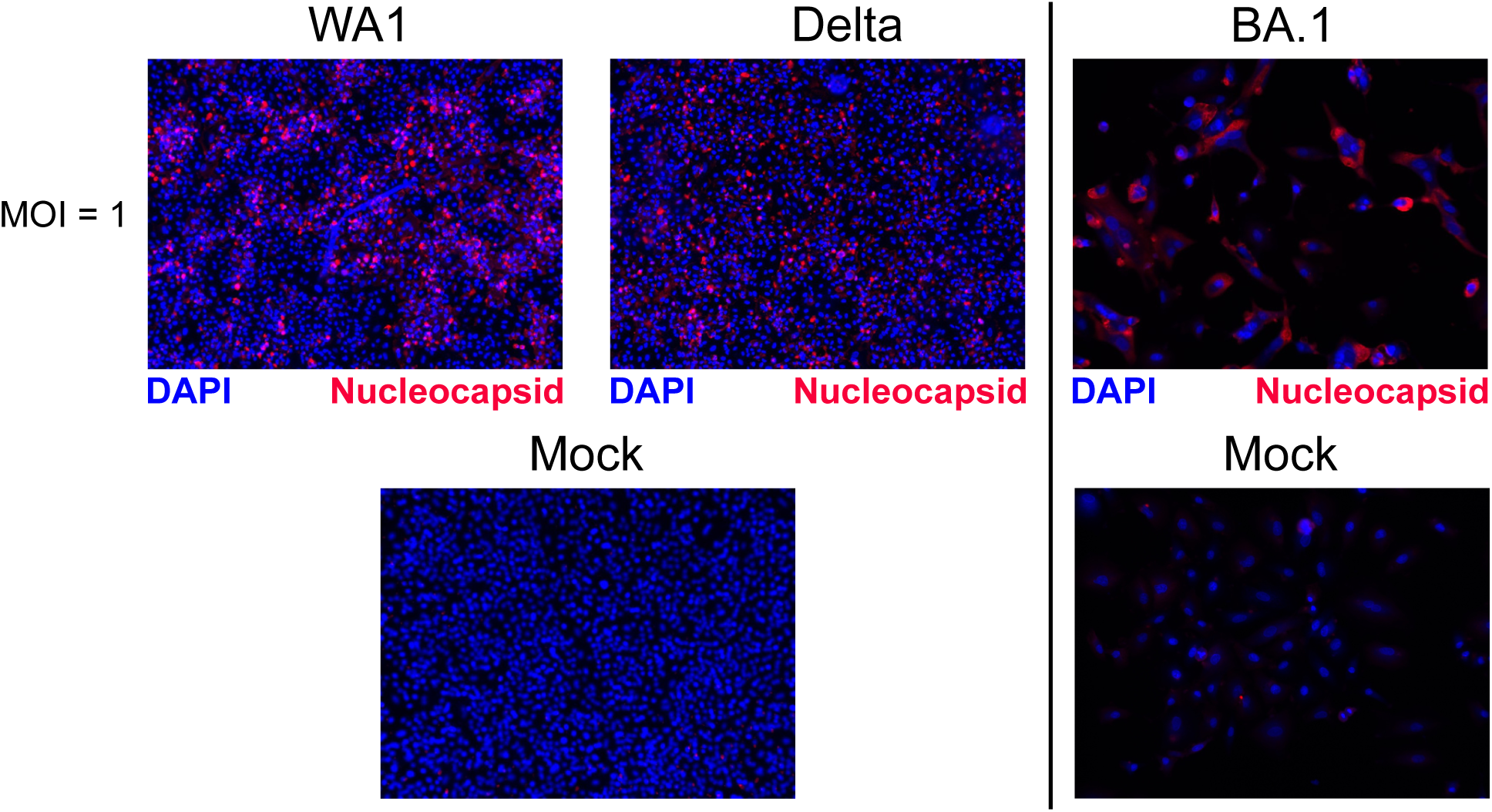
Infection efficiency of variants of SARS-CoV-2 in A549-ACE2 cells. Infection is assessed by immunofluorescence against the nucleocapsid protein, in comparison to cellular nuclei, stained by DAPI.

**Supplementary Figure 3.**
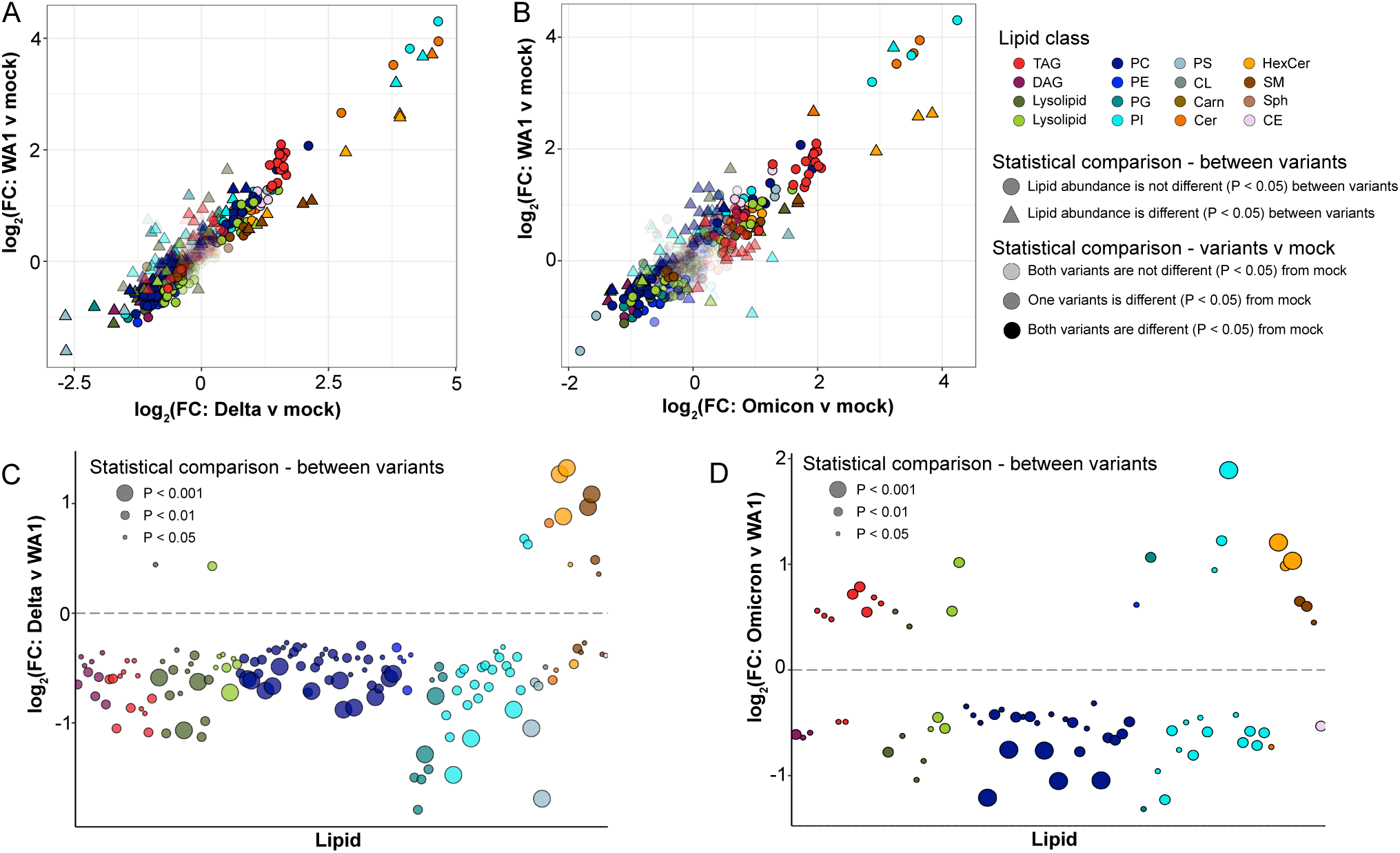
Lipidomics of variants of SARS-CoV-2 (**A**) log2FC of WA1 v mock compared to log2FC of delta v mock. (**B**) log2FC of WA1 v mock compared to log2FC of omicron v mock. Abbreviations: TAG = triacylglycerol; DAG = diacylglycerol; PC = phosphatidylcholine; PE = phosphatidylethanolamine; PG = phosphatidylglycerol; PI = phosphatidylinositol; PS = phosphatidylserine; CL = cardiolipin; carn = acylcarintine; Cer = ceramide; HexCer = hexosylceramide; SM = sphingomyelin; Sph = sphingosine; CE = cholesterol ester. (**C**) log2FC of Delta v WA1 for individual lipid species; only significantly changed (ANOVA, Tukey adjusted P < 0.05) lipid species are shown. (D) log2FC of Omicron v WA1 for individual lipid species; only significantly changed (ANOVA, Tukey adjusted P < 0.05) lipid species are shown.

